# Engineering novel AAV capsids by global de-targeting and subsequent muscle-specific tropism in mice and NHPs

**DOI:** 10.1101/2025.05.19.654800

**Authors:** Yue Pan, Yujian Zhong, Huan Chen, Youwei Zhang, Zhiyong Dai, Junlin Chen, Keqin Tan, Xiaoqu Chen, Danlan Qiu, Longxiang Sheng, Xinping Tan, Ying Fan, Ye Bu, Zexin Zhou, Zhiming Yang, Rui Duan, Min Guan, Guangping Gao, Huapeng Li

## Abstract

Recombinant adeno-associated viral (rAAV) vectors are a potent tool, but their clinical application is restricted by insufficient target tissue transduction and liver toxicity. We employed a novel two-step engineering strategy to create novel rAAV capsids with global tissue de-targeting, then produced strong tissue-specific expression by adding a peptide sequence. We created a novel capsid, AAV.Zero1, with globally de-targeted transduction by loop swapping domains from AAV9 into AAV2. Making an R585A substitution (AAV.Zero2) re-targeted tissues but deleting residues 585–587 (AAV.Zero3) abrogated transduction. Inserting a myogenic peptide into AAV.Zero3 produced a novel capsid (AAV.eM) with strong muscle-specific transgene expression while maintaining minimal off-target expression, including in liver, which was conserved in two mouse strains and non-human primates. AAV.eM showed similar expression as the leading myotropic vector MyoAAV.4A but had a more favorable safety profile. Importantly, AAV.eM was able to functionally rescue a mouse model of Duchenne Muscular Dystrophy following systemic delivery of a micro-dystrophin gene. Thus, AAV.eM is an improved myotropic rAAV capsid that de-targets other tissues, especially the liver, and proof-of-concept for a platform to create capsids with specific properties that translate across species by addition of peptides onto low transduction backbones.

## Introduction

Recombinant adeno-associated virus (rAAV) vectors are the most commonly used viral platforms for gene delivery[1–4]. However, two particular challenges limit their use in clinical practice: many of the delivered capsids are sequestered in the liver after systemic administration and there is insufficient transduction of intended target organs and cells to achieve therapeutic effects[5]. High viral doses (∼1–3E14 vg/kg) are usually required for the treatment of muscle-associated diseases to achieve optimal therapeutic outcomes, which presents challenges for virus production as well as concerns for liver toxicity and other host anti-AAV immune responses[6, 7]. Many clinical programs employ systemic administration of AAV to target non-hepatic tissues, such as muscle (*e.g.*, for muscular dystrophies; clinicaltrials.gov NCT03375164, NCT03368742, and NCT03362502). However, the liver is the primary organ that AAV particles transduce, which can cause toxicity and complicate the treatment. High dose systemic AAV administration is known to carry significant safety risks and was recently implicated as a contributing factor to the death of at least five patients in trials for X-linked myotubular myopathy[8–10]. Thus, strategies to de-target AAV transduction of the liver has been an area of active interest in the gene therapy field[11, 12].

Capsid engineering to optimize tropism for specific cells and tissues is a promising strategy to bolster the clinical efficacy of AAV vectors by reducing the viral dosage needed to achieve therapeutic effects[13, 14]. Using less vector to achieve therapeutic outcomes will help improve safety, overcome pre-existing immunity to natural AAV serotypes, lower the risk of host immune responses, and decrease manufacturing costs. AAV tropism is determined by the receptor-binding motifs of its capsid, which facilitates viral attachment to the surface of cells and subsequent cellular entry. Therefore, deciphering the structure-function relationship of these motifs and engineering specific capsid dynamics—from the nuanced receptor-ligand interactions to post-attachment processes such as internalization, sorting, endosomal escape, nuclear import, uncoating and vector genome processing—could allow for AAV variants to have tailored tropisms[15].

Effective viral transduction culminates in transgene expression, and so detection of intact virus capsid or vector genome is not sufficient to confirm successful transduction. Consequently, transcription-oriented screening platforms, including TRACER and DELIVER, have emerged as potent tools in novel capsid variant development[16, 17]. While bioinformatics platforms are being used to improve capsid protein engineering, *in vivo* validation remains necessary for identifying potent and target tissue-enhanced capsids. Intra-species and -strain differences in transgene expression following AAV transduction present another challenge, as numerous AAV capsids that were engineered to have enhanced transduction efficiency and/or specific tropism in murine models have failed to replicate their properties in non-human primate (NHP) studies[18–22].

In this study, we present a synergistic bioinformatic and rationally designed experimental approach in the development of three novel capsids. AAV.Zero1 and AAV.Zero3 have broadly low levels of transduction in diverse mouse tissues which can be used as universal backbones for engineering targeted tissue tropism through the addition of peptides with known properties. As proof-of-principle for this strategy, we incorporated a muscle-targeting peptide into AAV.Zero3 to yield a novel capsid, AAV.eM, which shows considerable promise for high gene transfer specifically to the musculature with minimal liver transduction following systemic administration in both rodents and NHPs.

These novel capsids were generated by rationally designed, capsid structure-guided, and bioinformatics-based loop swapping, followed by myotropic peptide insertion. They demonstrated similar transduction profiles in both murine and NHP models (*i.e.*, all showed reduced liver and AAV.eM showed enhanced muscle tropism). We suggest that this approach can be applied to create many additional capsids to target a variety of tissues with cross-species tropism. This rational design approach has the potential to improve the efficiency of capsid design as well as generate more tissue- and cell type-specific targeting for therapeutic and safe gene delivery while reducing toxicities caused by expression in off-target tissues, especially the liver. Improved vector design and efficacy will reduce the viral dose that needs to be delivered and the amount of vector that needs to be manufactured for both pre-clinical and clinical development as well as accelerate the development of optimal AAV gene therapies for numerous human diseases.

## Results

### Chimeric capsid variants exhibit significantly reduced hepato-tropism in mouse

Numerous AAV serotypes, including natural variants AAV7, AAV8, and AAV9 as well as novel engineered variants, exhibit high liver tropism following intravenous administration[1, 2]. To limit hepatic and systemic toxicities, we sought to engineer capsids that de-target the liver by modifying motifs of the AAV2 capsid, which has reduced liver tropism compared to AAV9. In comparing the 3D structure of AAV2 (PDB: 6IH9) and AAV9 (PDB: 3UX1) capsids, we noted that the surface-exposed loop IV is the farthest protrusion of the viral capsid in the 3-fold axis of symmetry. Binding interactions with cell-surface receptors occur near this axis of the AAV9 capsid, where loops IV and VIII form neighboring surface loops within a single monomer and form into three adjacent surface loops with loop V from anther monomer[23]. We noted that the positions of loops IV, V, and VIII in AAV2 show a similar pattern as AAV9 (Fig. 1A). Loop IV has been reported to be critical for the efficient liver transduction of AAV8[24] and has previously been implicated in neutralizing antibody binding[25]. We hypothesized that there is a structure-function relationship between these domains and their host-interacting properties which generate structural context-dependent tissue tropisms, so we introduced amino acid sequence diversity into loops IV and V. We replaced two loops of AAV2 (IV loop: R447–Q461 and V loop: K490–D494) with corresponding AAV9 sequences (IV loop: K449–K462 and V loop: T491–Q495), resulting in the chimeric capsid AAV.Zero1. The R585 amino acid is the key residue mediating binding of AAV2 to heparin sulfate proteoglycan (HSPG)[26], which is the major host cell receptor responsible for AAV2 hepato-tropism. Thus, we introduced additional mutations into the AAV.Zero1 capsid sequence which we hypothesized would further reduce liver tropism, creating chimeric serotypes AAV.Zero2 (R585A) and AAV.Zero3 (R585GN deletion) (Fig. 1B). We assessed the transduction profiles of these three new capsid variants by assessing their efficiency for transducing five immortalized cell lines from different tissues, kidney HEK293T, myoblast C2C12, hepatoma Huh-7, cervical cancer cell HeLaRC32, and hamster ovary CHO-K1, and found that they were distinct from AAV2 (Supplementary Fig. 1A). The expression of the delivered EGFP transgene was assessed by quantifying the fluorescence intensity from imaging of HEK 293T and C2C12 cells (Supplementary Fig. 1B). The analysis of relative vector genome copy number within cells was used to determine how efficient these AAV serotypes were at entering each of the cell types (Supplementary Fig. 1C), which generally correlated with the EGFP expression (compare Supplementary Fig. 1B and C).

**Figure 1.**
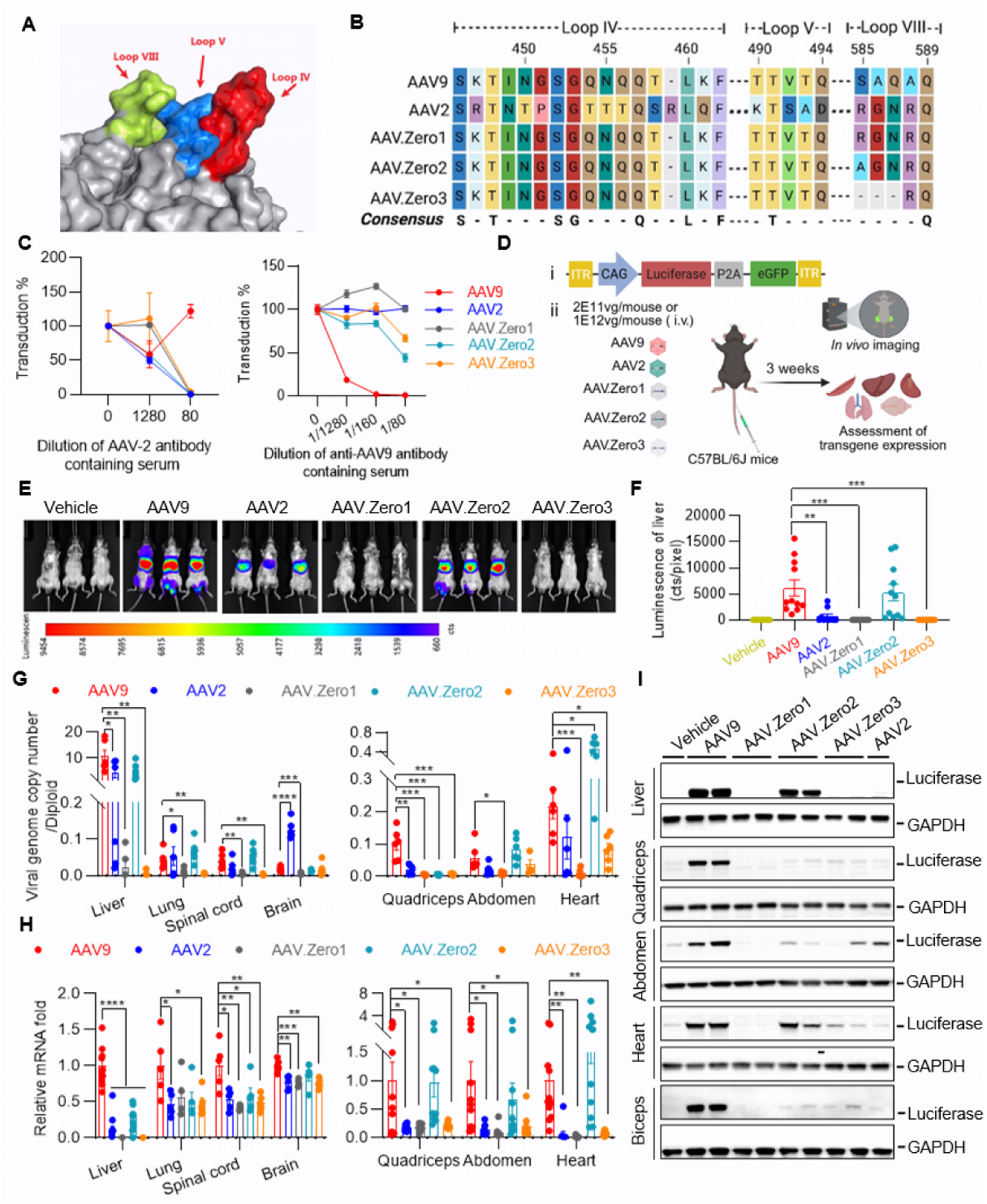
Chimeric capsid variants exhibit dramatical reductions in liver transduction. A. Capsid spike formed by loops IV and VIII of one AAV2 monomer interacting with the loop V of a second monomer. B. Sequence alignments of the variable regions of AAV.Zero1, AAV.Zero2, and AAV.Zero3 compared with AAV2 and AAV9. C. Different dilutions of AAV2 (left) or AAV9 (right) antibody-containing serum were incubated with constant amounts of AAV serotypes AAV9-, AAV2-, AAV.Zero1-, AAV.Zero2-, or AAV.Zero3-CAG-Fluc-P2A-EGFP at MOI=1E6 and tested for neutralization of transduction in an *in vitro* assay. Data are presented as mean ± SEM (n=3–6). D. Schematic illustration of virus administration and biodistribution assessment process. (i) CAG-luciferase-P2A-eGFP was packaged as reporter transgene. (ii) Experimental workflow in mice. E–I. 8-week-old C57BL/6J mice systemically injected with 2E11 vg per mouse (∼8E12 vg/kg) of AAV9-, AAV2-, AAV.Zero1-, AAV.Zero2-, or AAV.Zero3-CAG-Fluc-P2A-EGFP and data was collected 21 days post-injection. Representative whole body *in vivo* bioluminescence (E) and quantification of firefly luciferase luminescence (F). The *p* value was calculated between AAV9 and each other group by unpaired Student’s *t*-test (n=10–11), ∗∗*p* < 0.01, ∗∗∗*p* < 0.001. G. Viral genome copy number per diploid from mice tissues detected by droplet digital PCR. The *p* values were calculated between AAV9 and each other group by unpaired Student’s *t*-test (n=6), ∗*p* < 0.05, ∗∗*p* < 0.01, ∗∗∗*p* < 0.001, ∗∗∗∗*p* < 0.0001. H. Quantification of fold difference in *Fluc* mRNA expression. The *p* values were calculated between AAV9 and each other capsid by unpaired Student’s *t*-test (n=10–11, except n=4–5 for lung, spinal cord, and brain), ∗*p* < 0.05, ∗∗*p* < 0.01, ∗∗∗*p* < 0.001, ∗∗∗∗*p* < 0.0001. I. Representative western blot images detecting luciferase and GAPDH of liver, quadriceps, abdomen, heart, and biceps.

The presence of pre-existing anti-AAV antibodies in human populations represents a major hurdle for *in vivo* gene therapy because the antibodies neutralize AAV-based gene therapy vectors[27, 28]. Therefore, we evaluated the cross-reactivity of anti-AAV2 and anti-AAV9 antibodies against the three newly developed capsids using a neutralization inhibition assay. These novel vectors are derived from the AAV2 serotype and the results indicate that they cross-reacted with antibodies produced against AAV2 (Fig. 1C).

We intravenously administered AAV9-, AAV2-, AAV.Zero1-, AAV.Zero2-, and AAV.Zero3-CAG-luciferase-P2A-EGFP to adult C57BL/6J mice at dosages of 2E11 (Fig. 1D–I) or 1E12 (Fig. 1D and Supplementary Fig. 1D–G) viral genomes (vg) per mouse to investigate the transduction profile and biodistribution of the chimeric variants compared to the parental capsids. The *in vivo* bioluminescence imaging data indicate that AAV.Zero1 and AAV.Zero3 had weaker luciferase activity in the liver compared to AAV9 (Fig. 1E, F and Supplementary Fig.1D, E). In agreement with these findings, AAV.Zero1 and AAV.Zero3 showed significantly fewer viral particles (Fig. 1G) and produced lower levels of luciferase mRNA transcripts (Fig. 1H, Supplementary Fig. 1F) in the liver compared to AAV9. Western blotting analysis similarly shows that AAV.Zero1 and AAV.Zero3 produced low levels of luciferase protein in the liver compared to the AAV9 (Fig. 1I and Supplementary Fig. 1G). We additionally detected the biodistribution patterns of the three new capsids in a selection of other tissues, including quadriceps, abdomen, heart, lung, spinal cord, brain, kidney, and eye. AAV.Zero1 and AAV.Zero3 transduced fewer viral particles and induced lower transgene expression in all tested organs at both low (Fig. 1G–I) and high (Supplementary Fig. 1F, G) viral dosages, suggesting that they can both be used as starting backbones with globally reduced tissue tropism to engineer capsids for specific organ tropism.

The capsid protein sequence of AAV.Zero1 is identical to the AAV2 capsid except for loops IV and V, which are the sequences for the corresponding loops in AAV9. This modification resulted in a broad loss of transduction in every organ tested for AAV.Zero1 compared to its parental capsid AAV2. Remarkably, making only one substitution in the AAV.Zero1 sequence (R585A) to generate AAV.Zero2 restored viral particle transduction and strong transgene mRNA expression in several muscular organs, including quadriceps, abdomen, heart (Fig. 1G, H, I and Supplementary Fig. 1F, G), and eye (Supplementary Fig. 1F). The difference in the biodistribution of transduced cells between AAV.Zero1 and AAV.Zero2 suggests that the R585 residue can functionally serve as an On/Off switch to transduce multiple tissues. AAV.Zero3 has a similar biodistribution profile to AAV.Zero1 of low transduction levels in all tested tissues, but does not contain the R585–N587 sequence, and so lacks the On switch.

### Enhanced muscle specificity achieved through the incorporation of RGD-containing peptides into a chimeric capsid

We next sought to engineer tissue-specific targeting properties into one of the low transduction capsids by the insertion of a peptide sequence. For enhanced muscle tropism, we integrated three previously characterized myotropic RGD-containing peptides[29], 2A, 4E, and 4A, into AAV.Zero3 at site 585, generating new capsids AAV.Zero3 2A, AAV.Zero3 4E, and AAV.eM, respectively (Fig. 2A). We performed *in vitro* cell transductions and found that these novel vectors showed detectable transduction of C2C12 cells, although notably weaker than AAV2 and MyoAAVs, but not HEK293T, Huh-7, HelaRC32, and CHO-K1 cells (Fig. 2B and Supplementary Fig. 2A). However, the AAV genome DNA copy numbers (a measure of cell entry efficiency) were not predictive of the EGFP fluorescence intensity that was generated (a measure of transgene expression) (compare Fig. 2B and Supplementary Fig. 2B). We next evaluated the cross-reactivity of these vectors by neutralization inhibition assays with anti-AAV2 and anti-AAV9 antibodies (Supplementary Fig. 2C, D). The capsids containing peptide insertions showed similar cross-reactivity as their parental backbone AAV.Zero3, with noticeable inhibition by antibodies produced against AAV2 and serologic profiles being more comparable to AAV2 than AAV9. As expected, these chimeric capsids were resistant to all concentrations of anti-AAV9 antibodies tested (Supplementary Fig. 2D).

**Figure 2.**
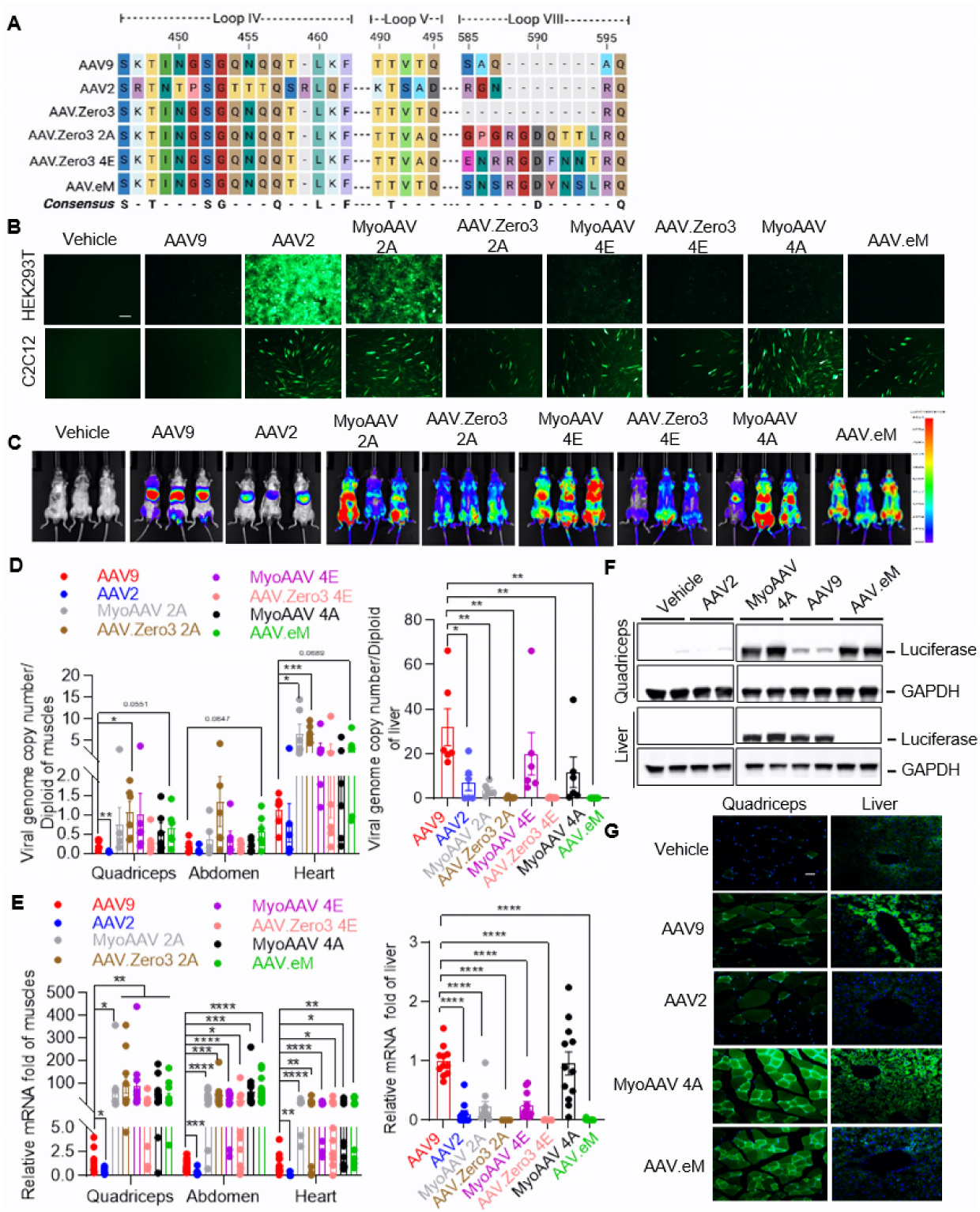
RGD-containing peptides enable chimeric capsid selectively target to muscle with high efficiency in C57BL/6J mice. A. Sequence alignments of the RGD-containing peptides 2A, 4E, and 4A inserted into AAV.Zero3 at R585. B. Representative images of *in vitro* transduction in HEK 293T and C2C12 cell lines were transduced at MOI=1E5 with AAV9-, AAV2-, MyoAAV 2A-, AAV.Zero3 2A-, MyoAAV 4E-, AAV.Zero3 4E-, MyoAAV 4A-, or AAV.eM-CAG-Fluc-P2A-EGFP 72 hours after transduction. Scale bar: 100 μm. C–E. 8-week-old C57BL/6J mice were systemically injected with 2E11 vg per mouse (∼8E12 vg/kg) AAV9-, AAV2-, MyoAAV 2A-, AAV.Zero3 2A-, MyoAAV 4E-, AAV.Zero3 4E-, MyoAAV 4A-, or AAV.eM-CAG-Fluc-P2A-EGFP. Representative whole body *in vivo* bioluminescence images 21 days post-injection (C). Viral genome copy number per diploid from mice tissues, including quadriceps, abdomen, and heart (left) and liver (right) detected by qPCR with standard curve (D). The *p* value was calculated between AAV9 and each other capsid by unpaired Student’s *t*-test (n=6), \**p* < 0.05, \*\**p* < 0.01, \*\*\**p* < 0.001. Quantification of the *Fluc* mRNA in mice tissues, including quadriceps, abdomen, and heart (left) and liver (right) 21 days post-injection (E). The *p* value was calculated between AAV9 and each other capsid by unpaired Student’s *t*-test (n=11–17), \**p* < 0.05, \*\**p* < 0.01, \*\*\**p* < 0.001, \*\*\**p* < 0.001. F–G. 8-week-old C57BL/6J mice were systemically injected with 2E11 vg per mouse (∼8E12 vg/kg) AAV9-, AAV2-, MyoAAV 4A-, or AAV.eM-CAG-Fluc-P2A-EGFP. Representative western blot images detecting luciferase and GAPDH of quadriceps and liver tissues 21 days post-injection (F). Representative cross-section fluorescent images of quadriceps and liver tissues 21 days post-injection (G). Green: EGFP, Blue: DAPI. Scale bar: 25 μm.

We intravenously injected adult C57BL/6J mice with parental capsid AAV9 or AAV2, novel chimeric capsid AAV.Zero3 2A, AAV.Zero3 4E, or AAV.eM, or established myotropic capsid MyoAAV 2A, MyoAAV 4E, or MyoAAV.4A vectors packaged with a CAG-luciferase-P2A-EGFP transgene cassette at a dose of 2E11 vg per mouse to investigate their transduction profiles and biodistributions *in vivo*. Three weeks post-injection, whole body and organ bioluminescence imaging shows that the novel AAV.Zero3 2A, AAV.Zero3 4E, and AAV.eM capsids produced comparable luciferase expressions in muscles relative to the previously developed myotropic MyoAAV 2A, MyoAAV 4E, and MyoAAV.4A capsids and notably more than the parental AAV2 and AAV9 capsids (Fig. 2C and Supplementary Fig. 2E). Similar results were observed for viral genome DNA (Fig. 2D), luciferase mRNA (Fig. 2E), and protein levels (Fig. 2F and Supplementary Fig. 2F) as well as fluorescence imaging of individual tissue sections (Fig. 2G) that the novel and established myotropic vectors showed high transgene expression across multiple muscle groups. Importantly, the novel vectors appeared to induce less transgene expression in the liver compared to the established myotropic vectors. These results suggest that the insertion of RGD-containing peptides can confer enhanced muscle transduction efficiency onto the globally de-targeted backbone AAV.Zero3 while maintaining its liver de-targeting properties (Fig. 2D–G and Supplementary Fig. 2F). Thus, this approach can be used to improve AAV capsid targeting of specific cells and tissues while de-targeting the liver.

### AAV.eM both efficiently and specifically targets muscle across rodent species

A major challenge for engineering tissue-specific tropism of a vector is replicating transduction profiles across species. We chose AAV.eM as a candidate for further development and cross-species validation because it contains the 4A peptide, which has previously been shown to have cross-species myo-trophic properties in capsids[30, 31]. Our initial experiments were in C57BL/6J mice, so we next intravenously injected adult BALB/c mice with AAV9-, AAV2-, MyoAAV.4A-, or AAV.eM-CAG-luciferase-P2A-EGFP at a dose of 2E11 vg per mouse (∼8E12 vg/kg). At 3 weeks post-injection, we analyzed *in vivo* luciferase activity and transgene expression in a variety of tissues (Fig. 3A). Whole-organ bioluminescence imaging shows that AAV.eM induced significantly higher luciferase expression in limbs compared to its parental AAV2 and AAV9 capsids and the previously developed myotropic vector MyoAAV.4A (Fig. 3B). Similarly, AAV.eM produced significantly higher viral genome copies per cell transduced (Fig. 3C) and luciferase mRNA expression (Fig. 3D) in skeletal muscles and heart compared to MyoAAV 4A. We additionally assayed the brain and kidney as other non-targeted tissues and found that they showed similar transduction profiles for AAV2, AAV9, MyoAAV 4A, and AAV.eM (Figure 3C, D). AAV.eM induced higher EGFP protein expression than AAV2, AAV9, and MyoAAV 4A in triceps, quadriceps, and heart muscles while expressing much lower levels in the liver (Fig. 3E). Collectively, these findings suggest that AAV.eM shows proficient muscle targeting and minimal liver targeting, providing proof-of-concept that incorporating a specific peptide into a low universal transduction backbone can be used to engineer capsids with specific tropism for desired tissues similarly in two mouse strains.

**Figure 3.**
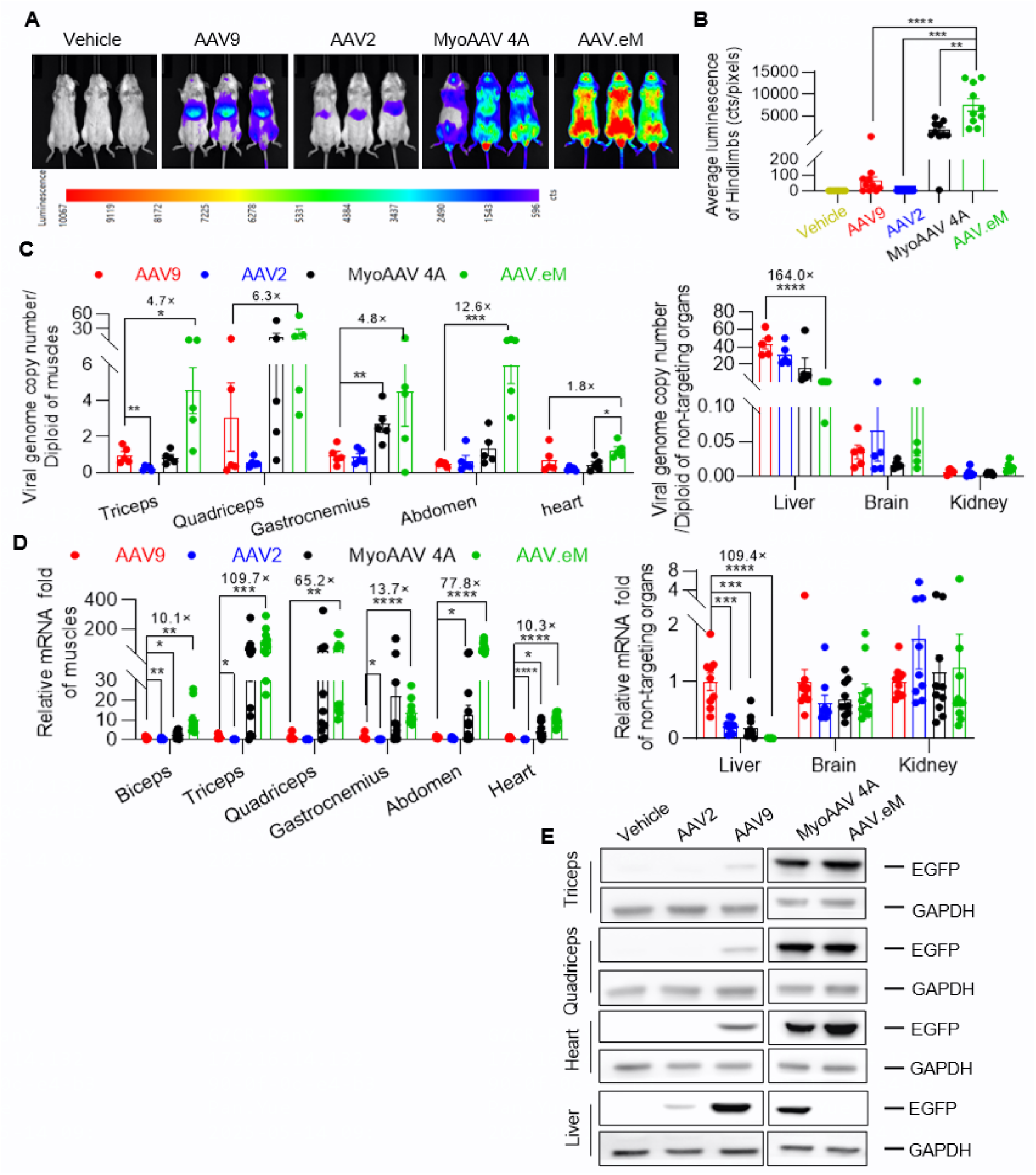
AAV.eM is both efficient and specific in muscle-targeting cross rodent species. 8-week-old BALB/c mice were systemically injected with 2E11 vg per mouse (∼8E12 vg/kg) of AAV9-, AAV2-, MyoAAV 4A-, or AAV.eM-CAG-Fluc-P2A-EGFP and data was collected 21 days post-injection. A. Representative whole body *in vivo* bioluminescence images. B. Quantification of firefly luciferase luminescence from hindlimbs. The *p* value was calculated between AAV.eM and other groups by Student’s *t*-test (n=7–10), \*\**p* < 0.01, \*\*\**p* < 0.001, \*\*\*\**p* < 0.0001. C. Viral genome copy number per diploid from mouse tissues detected by qPCR using a standard curve. The *p* value was calculated between AAV9 and each other groups by Student’s *t*-test (n=5), \**p* < 0.05, \*\*\**p* < 0.001, \*\*\*\**p* < 0.0001. D. Quantification of the *Fluc* mRNA in biceps, triceps, quadriceps, gastrocnemius, abdomen, and heart muscles as well as the liver, brain, and kidney. The *p* value was calculated between AAV9 and each other groups by Student’s *t*-test (n=6–10), \**p* < 0.05, \*\**p* < 0.01, \*\*\**p* < 0.001, \*\*\*\**p* < 0.0001. E. Representative western blot images detecting EGFP and GAPDH of triceps, quadriceps, heart, and liver.

### AAV.eM shows high muscle transduction and low liver toxicity in non-human primates

There is a pressing demand to extrapolate findings from rodent studies to human disease applications, and NHPs serve as invaluable models in this translation from pre-clinical investigation to clinical trials[32, 33]. Given that capsids engineered for tissue-specific tropism in mice have been shown to not universally translate to NHPs[34–37], we examined the transduction profile of our novel AAV.eM vector in primates. We intravenously injected adult macaques (*Macaca fascicularis*) with MyoAAV.4A- and AAV.eM-CAG-luciferase-P2A-EGFP at a moderate dosage of 3E13 vg/kg. After AAV injection, hepatotoxicity was monitored over two weeks by measuring circulating alanine aminotransferase (ALT) and aspartate aminotransferase (AST) levels, two reliable markers of liver injury. We found that ALT and AST levels after delivery of MyoAAV.4A peaked on the third day post-injection, reaching 17.9× (AST) and 13.2× (ALT) higher than the pre-injection levels (Fig. 4A). However, in the animals that received AAV.eM, the serum AST and ALT levels remained consistently similar to the level measured before virus injection throughout the monitoring phase, suggesting a superior safety profile post-systemic administration. We assessed muscle damage over 14 days after virus injection as measured by circulating creatine kinase (CK) levels and found that MyoAAV.4A and AAV.eM did not cause notable changes in serum CK relative to the level before injection (Fig. 4A).

**Figure 4.**
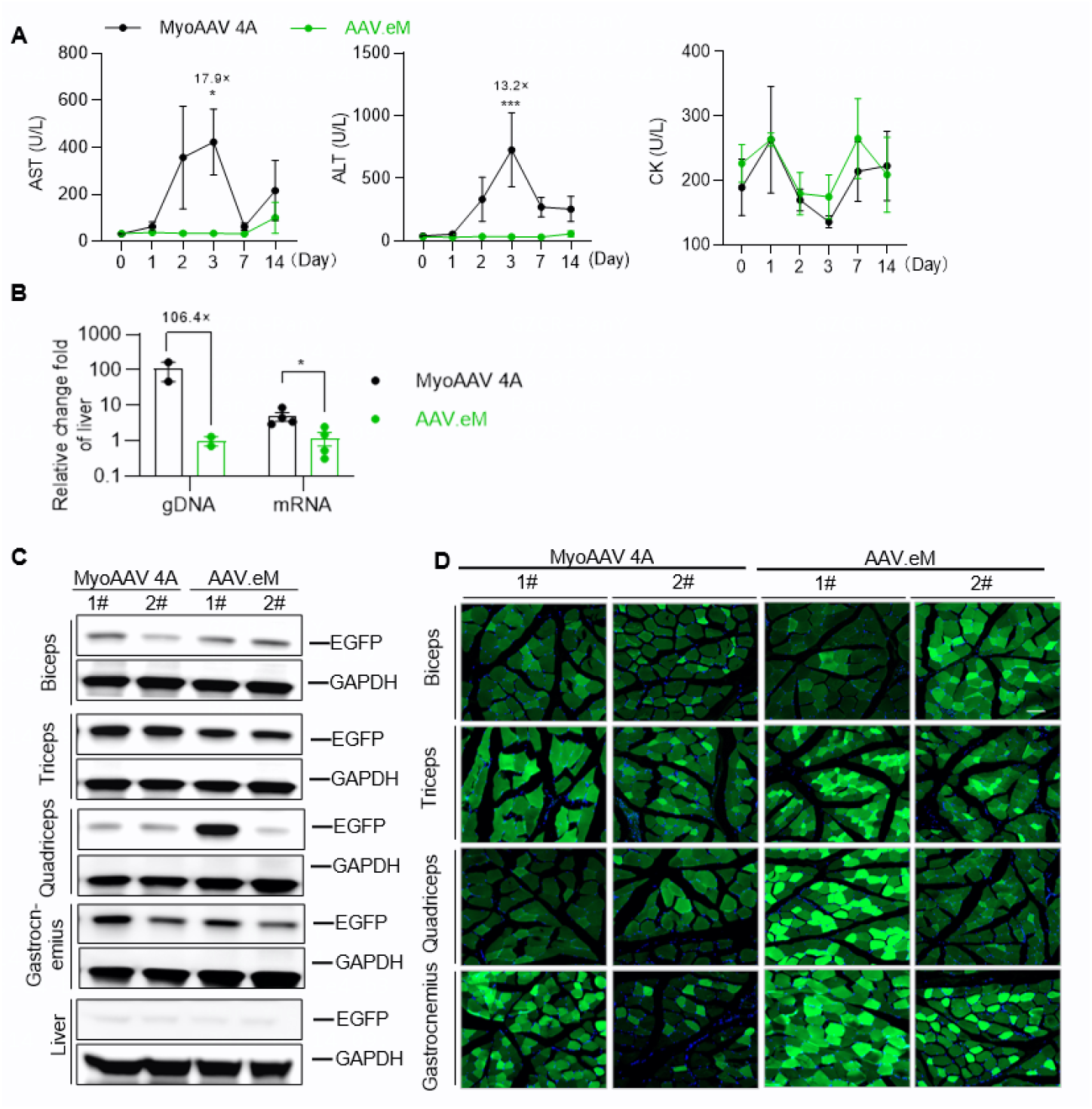
AAV.eM causes less liver toxicity without sacrificing the transduction efficiency in non-human primates. NHPs were given intravenous administration of 3E13 vg/kg of AAV.eM- or MyoAAV 4A-CAG-Fluc-P2A-EGFP. A. Hepatic enzyme (AST and ALT) and creatine kinase (CK) levels were monitored prior to and after dosing. The *p* values were calculated by two-way ANOVA with Tukey-Kramer multiple comparisons test. (n=5 for AST and ALT, n=3 for CK), \**p* < 0.05, \*\*\**p* < 0.001. AST, aspartate aminotransferase; ALT, alanine aminotransferase. B. Quantification of relative vector genome per diploid genome and *Fluc* mRNA relative fold-change in the liver of NHPs 28 days post-injection. The *p* value was calculated by unpaired Student’s *t*-test (n=4 for mRNA and n=2 for gDNA), \**p* < 0.05. C. Representative western blot image detecting EGFP and GAPDH of biceps, triceps, quadriceps, gastrocnemius, and liver sampled 14 days post-injection. D. Representative cross-section fluorescent images of biceps, triceps, quadriceps, and gastrocnemius sampled 28 days post-injection. Green: EGFP, blue: DAPI, Scale bar: 100 μm. 1# and 2# represents two independent monkeys of each group.

As part of the transduction process, the packaged AAV genome undergoes conversion from single stranded DNA (ssDNA) into double stranded DNA (dsDNA) before the transgene is expressed[38]. We assessed the abundance of vector genomes in the livers of the monkeys and found that AAV.eM showed less liver transduction and less transgene expression than MyoAAV.4A (Fig. 4B). We also assessed various muscle tissues, including biceps, triceps, quadriceps, and gastrocnemius, and found that AAV.eM produced a comparable number of vector genome copies and transgene expression in these tissues as MyoAAV.4A on day 28 after virus injection (Supplementary Fig. 3A). In agreement with this, western blotting analysis also indicated that AAV.eM expressed the EGFP protein in different muscle tissues at levels similar to MyoAAV 4A and both showed little EGFP expression in the livers (Fig. 4C). The transgene mRNA (Fig. 4B and supplementary Fig.3A) and protein (Fig 4C) levels were consistent with EGFP fluorescence imaging results of the muscle sections (Fig. 4D). To comprehensively characterize the tissue distribution and transgene expression profiles of the two capsids in NHPs, we quantified viral genome copy numbers across 17 distinct tissue types, encompassing both muscle tissues (including biceps, triceps, quadriceps, gastrocnemius, tibialis anterior, soleus, abdominal wall, diaphragm, and heart) and organs (liver, lung, brain, kidney, and spleen) (Supplementary Fig. 3B). AAV.eM showed comparable transduction as MyoAAV.4A in trunk muscles and cardiac tissues, but notably less transduction in the liver (Supplementary Fig. 3B). Gene expression analysis revealed AAV.eM and MyoAAV.4A exhibited broadly comparable transgene mRNA expression across most tissues, with the notable exception of the liver and brain, where the novel AAV.eM capsid produced expression levels that were approximately two orders of magnitude lower than the established MyoAAV.4A capsid (Supplementary Fig. 3C). Overall, these findings indicate superior liver de-targeting of AAV.eM compared to the previously engineered muscle-tropic vector MyoAAV.4A while maintaining similar muscle-targeting efficiency in NHPs.

### A low-background capsid backbone is amenable to a variety of peptides directing tissue-specific transduction with minimal off-target effects

We considered the possibility that our globally de-targeted capsids could be specific to the MyoAAV peptides and therefore not broadly useful for producing tissue-specific tropism. To evaluate the generalizability of our low-background AAV.Zero3 capsid for peptide-mediated tissue-specific targeting, we engineered additional variants by incorporating an RGD-containing peptide motif derived from TGF-β[39, 40], generating a new capsid designated PG016, and introduced peptides selected from a previously constructed RGD-containing randomized library, generating two additional capsids, PG017 and PG018 (Supplementary Fig. 4A). All three novel capsids demonstrated muscle-targeting efficiency comparable to that of MyoAAV 4A in murine models while exhibiting markedly reduced transduction in the liver, brain, and kidney (Supplementary Fig. 4B–D). Consistent results were observed in NHPs, where liver transduction was also significantly diminished when viral vectors were mixed in equal proportions and administered into monkeys via systemic injection (Supplementary Table 1). These findings indicate that all of the engineered variants preserved muscle tropism while minimizing off-target transduction in both mice and NHPs, supporting the broad applicability of the AAV.Zero3 backbone for peptide-guided, tissue-specific gene delivery.

### Systemic delivery of micro-dystrophin with AAV.eM produces functional correction in a Duchenne muscular dystrophy mouse model

To investigate the feasibility of using AAV.eM for *in vivo* delivery of therapeutic transgenes, we compared its delivery efficiency to AAV9, which is the delivery vector that has been used in clinical trials for Duchenne muscular dystrophy (DMD) (NCT03362502 and NCT03368742)[41, 42]. We systemically administered AAV9 or AAV.eM containing a truncated function-complementing dystrophin (micro-dystrophin, μDys) transgene expressed by the muscle-specific *MHCK7* promoter to the dystrophin-deficient B10 DMD-KO mouse model (4 bp deletion in *Dmd* exon 4)[43]. At 14 weeks post-injection, the whole-limb grip strength assay indicates that the muscle strength of AAV.eM-μDys-injected mice improved compared to both DMD-KO vehicle- and AAV9-injected animals, indicating functional rescue of muscle function (Fig. 5A). At 20 weeks post-injection, the serum CK level detected in AAV.eM-μDys-treated mice was significantly reduced compared to DMD-KO vehicle-treated group, indicating partial rescue of the severe muscle damage (Fig. 5B). We assessed biodistribution by quantifying viral genome copy number in various tissues and found that the AAV.eM-μDys-injected group showed higher μDys transgene delivery to multiple muscles, including biceps, triceps, quadriceps, gastrocnemius, abdomen and heart, compared to AAV9-μDys-injected animals, but less delivery to other organ tissues, including liver, lung, brain, and kidney (Fig. 5C). Similarly, we found that μDys mRNA levels were higher in muscles and lower in non-muscle tissues of mice injected with AAV.eM-μDys compared to AAV9-μDys (Fig. 5D). We assessed the expression of μDys protein levels in muscle tissues by immunofluorescence staining and found remarkably better restoration of dystrophin expression in biceps, triceps, quadriceps, and gastrocnemius muscles of mice treated with the AAV.eM-μDys vector compared to the AAV9-μDys vector, indicating higher targeted muscle transduction and lower off-target transduction (Fig. 5E). These data establish that AAV.eM-μDys shows therapeutic efficacy in treating DMD-KO mice when delivered by systematic administration which is superior to AAV9.μDys. This rationally engineered capsid produces enhanced muscle performance, reduced muscle damage, and elevated dystrophin mRNA and protein expression compared to the conventional AAV9 capsid, which are therapeutic targets that clinical trials are attempting to achieve in DMD patients. Thus, this vector and the design strategy used to produce it have the potential to deliver improved performance, reduced off-target transduction which can cause adverse events, and achieve therapeutic efficacy with lower dosages to improve the safety profile and reduce manufacturing costs.

**Figure 5.**
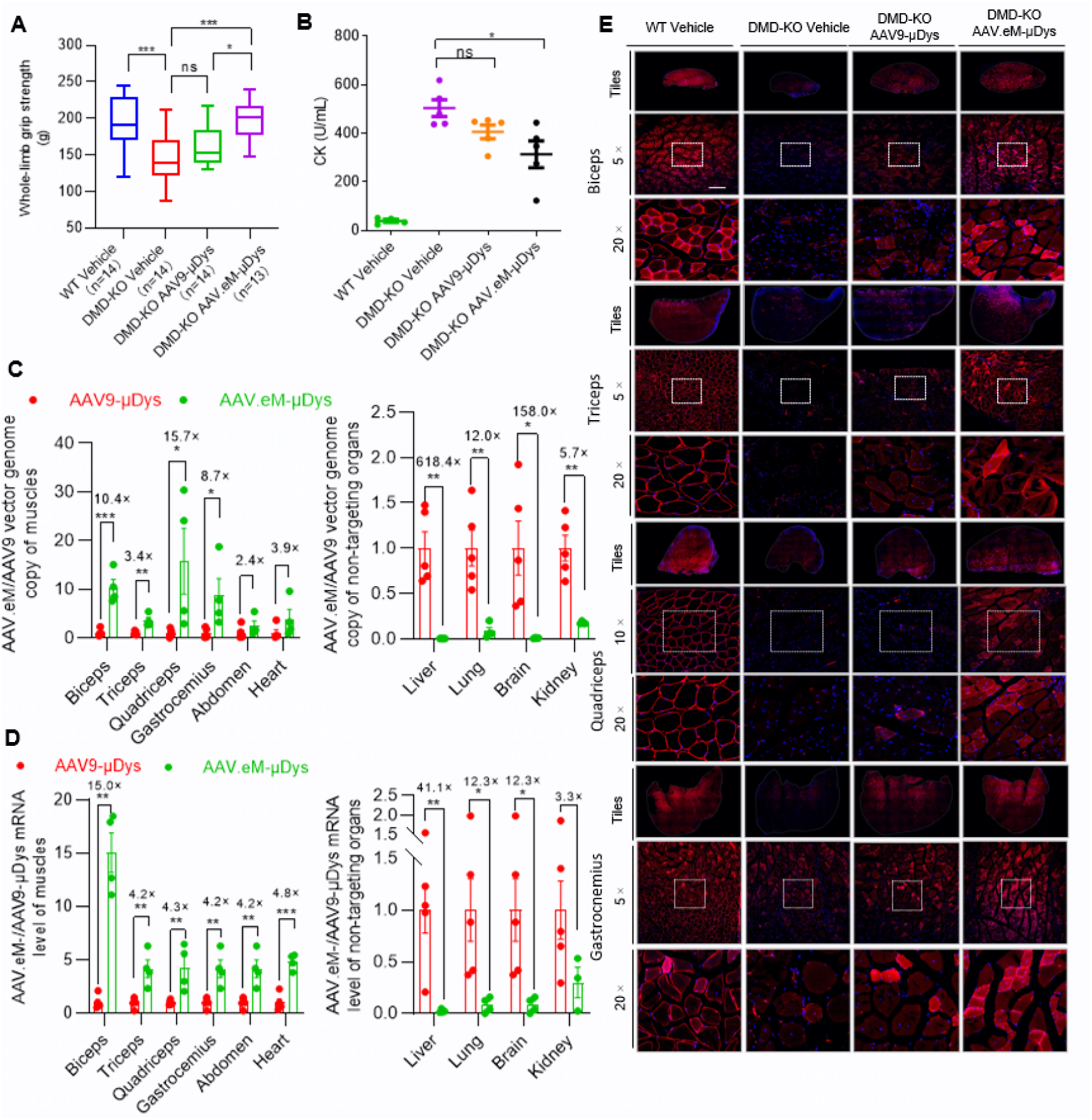
Systemic injection of AAV.eM-MHCK7-microdystrophin results in widespread micro-dystrophin expression and effective restoration of muscle function in adult DMD mice. 8-week-old wild-type B10 mice and B10 DMD-KO mice were systemically injected with vehicle or 2E13vg/kg of either AAV9 or AAV.eM-MHCK7-microdystrophin (μDys). A–B. The grip strength (A) and serum creatine kinase (CK) levels (B) were assessed 14 weeks post-injection. The *p* values were calculated by one-way ANOVA with Tukey-Kramer multiple comparisons test (n=13–14 for grip strength, n=4–5 for CK concentration). \**p* < 0.05, \*\*\**p* < 0.001, ns: not significant. C–E. Quantification of relative vector genome copy number (C), mRNA expression (D), and representative fluorescence images of cross-sections stained (E) for μDys 20 weeks post-injection. The AAV.eM-μDys expression data is normalized to the relative expression levels of AAV9-μDys. The *p* values were calculated by unpaired Student’s t-test (n=5 for AAV9-μDys, n=4 for AAV.eM-μDys except for kidney which is n=3), \**p* < 0.05, ** <0.01, \*\*\**p* < 0.001. Red: μDys, Blue: DAPI, Scale bar: 200 μm.

## Discussion

Recombinant AAV vectors represent a leading platform for gene therapy in multiple organs to treat numerous human diseases[44, 45]. Ideally, an AAV vector should robustly express the therapeutic transgene in the desired tissue with specificity and efficiency with a single systemic delivery at a safe dose. The development of effective AAV vector platforms in the gene therapy field has resulted from the synergy of efforts to improve capsid properties and vector genome design. The most common strategy to identify novel capsids with desired properties has been to screen a high-throughput library of capsids with random peptides inserted for directed evolution by a primary selection criterion (*e.g.*, enhanced targeting to an organ), followed by more focused subsequent screenings of candidates to compare the properties of each AAV capsid directly in animals[46–48]. However, this approach is expensive and labor-intensive, and only produces incremental improvements in capsid performance with no guarantee that the capsid properties will translate across species. In contrast, rational design approaches have been undertaken based on the knowledge of capsid structures and mechanisms of AAV binding to cell surface receptors, internalization, and cellular trafficking. Rational design approaches primarily seek to alter capsid properties by engrafting designed peptides onto its surface[49, 50]. In this study, we employed rational design principles to engineer novel capsids, starting with domain swapping two loops from AAV9 into AAV2 (creating AAV.Zero1) and alteration of the critical R585 residue (AAV.Zero2 containing an R585A substitution and AAV.Zero3 containing an R585GN deletion). AAV.Zero1 and AAV.Zero3 showed broadly lowered levels of transduction in major tissues compared to their parental capsids and specifically de-targeted the liver in two mouse strains, while AAV.Zero2 re-targets tissues with a single amino acid substitution. We then designed the AAV.Zero3 2A, AAV.Zero3 4E, and AAV.eM capsids to be muscle-tropic by inserting RGD-motif containing peptides at the R585 in the variable loop VIII of AAV.Zero3[51, 52]. These showed broadly similar profiles to the established myotropic vectors that the inserted peptides were identified from, so we examined the biodistribution of AAV.eM in NHPs because its inserted peptide has previously been shown to translate across species. We compared it to the established myotropic AAV vector MyoAAV 4A and found that the two capsids generally displayed comparable transduction efficiencies in various muscles of C57BL/6J mice, including limbs, abdomen, and heart. This distribution pattern and high level of expression in muscle tissues, as measured by *in vivo* bioluminescence as well as DNA, mRNA and protein levels, was replicated in the BALB/c mouse strain and NHPs, indicating robust and cross species-translatable muscle targeting properties. Importantly, the novel AAV.eM capsid showed a superior safety profile in NHPs compared to the established MyoAAV 4A capsid and functional correction delivering µDys in a mouse model of DMD. Thus, the AAV.eM capsid shows promise as a myotropic vector for systemic delivery of therapies to treat diseases that affect the musculature system or to achieve high serum levels of a protein using the muscles as an organ system for biomanufacturing. Importantly, AAV.eM showed minimal liver transduction and did not induce increases in levels of circulating AST and ALT in NHPs, which are indicators of hepatotoxicity. In AAV gene therapy clinical trials, adverse events related to hepatotoxicity have been widely reported[10, 42], with increased incidents at higher vector doses. Thus, the AAV.eM and similarly engineered vectors can be employed to more efficiently deliver gene therapies to target tissues to achieve therapeutic efficacy without inducing liver damage. Additionally, the enhanced gene expression efficiency of this vector can allow for lower viral doses to be administered to achieve therapeutic effects, which is beneficial for both safety risks and manufacturing costs. In addition to its *in vivo* properties, AAV.eM also exhibits enhanced biophysical stability and reduced aggregation relative to MyoAAV 4A during vector production (Supplementary Table 2). Notably, MyoAAV 4A displays a threshold-dependent instability, with a marked increase in aggregation observed at concentrations exceeding 1E13 vg/mL. In contrast, AAV.eM retains a monodisperse profile even at ultra-high vector genome titers, underscoring its superior suitability for large-scale manufacturing applications.

We attribute the de-targeting property of AAV.eM for non-muscle tissues to its backbone, AAV.Zero3, which showed broadly low-level transduction in major tissues, especially liver, after systematic delivery. Compared to AAV2, AAV.Zero3 and AAV.Zero1 showed very little liver transduction, suggesting that the loop IV and loop V domains are important in determining the liver tropism of AAV2, which was suggested in the report of the AAV.CAP-B10 capsid[53]. However, these two loop regions are not the only elements affecting hepato-tropism. The single R585A substitution to create AAV.Zero2 from AAV.Zero1 restores liver transduction, emphasizing the role of R585 in HSPG-binding and hepato-tropism that was previously reported[13]. Making H527Y and R533S substitutions in the AAV9 sequence was also reported to reduce liver transduction and making an N271D substitution in the sequence of AAV8 was reported to result in a liver de-targeting phenotype[54], suggesting that there may be several mechanisms that affect liver tropism[55].

These novel capsids with broad low transduction efficiency levels for tissues allow for rational design principles such as peptide insertion and domain swapping to be applied in order to create novel capsids with specific properties such as enhanced transduction efficiency for specific tissues or cells and reduced off-target expression in unintended tissues. We used this design principle to create three additional novel muscular-targeting capsids (PG016, PG017, and PG018) by insertion of RGD motif-containing peptides with sequence variety into the global de-targeting capsid backbone AAV.Zero3 to again produce muscle-specific tropism. These three capsid variants showed comparable muscle targeting as MyoAAV 4A and significantly lower transduction in the livers of mice and NHPs, suggesting that this engineering strategy to produce high tissue-specific expression with low off-target transduction that is conserved across species has broad applicability. Thus, this approach shows great potential to improve clinical therapeutic efficacy and safety profiles of rAAV capsids for gene therapies.

## Materials and Methods

### Construction of the capsid variants

The capsid sequences of AAV.Zero1, AAV.Zero2, and AAV.Zero3 were constructed by Gibson assembly (New England Biolabs (NEB), E5510S). The ∼5,000 bp plasmid backbone was recovered from restriction digestion of *SmiI* and *BshT* of pAAV2-RC. Fragments A to F were amplified from the AAV2 Rep-Cap plasmid template with primers containing the replaced sequence of AAV9 (primer sequences are list in Supplementary Table 3) (Takara Bio, R050Q). Primer pairs of each fragment are Cap-F/YJ69-R for A, YJ69-F/YJ72-R for B, YJ72-F/Cap-R for C, YJ72-F/247-R for D, 247-F/Cap-R for E, YJ72-F/248-R for F, and 248-F/Cap-R for G. The full sequence of AAV.Zero1 was assembled with fragments A, B, C, and plasmid backbone. The full sequence of AAV.Zero2 was assembled with fragments A, B, D, E, and plasmid backbone. The full sequence of AAV.Zero3 was assembled with fragments A, B, F, G, and plasmid backbone.

The peptide insertion variants were constructed in the same way as described above, with the differences being the PCR template (AAV.Zero3) and primers (see Supplementary Table 3). Primer pairs of each fragment are Cap-F/249-R for H, 249-R/Cap-R for I, Cap-F/250-R for J, 250-R/Cap-R for K, and Cap-F/YJ107-R for L, and YJ107-R/Cap-R for M. The full sequence of AAV.Zero3 2A was assembled with fragments H, I, and plasmid backbone. The full sequence of AAV.Zero3 4E was assembled with fragments J, K, and plasmid backbone. The full sequence of AAV.Zero3 4A was assembled with fragments L, M, and plasmid backbone.

### Cell lines

The human embryonic kidney cell line HEK 293T (American Type Culture Collection (ATCC), CRL-11268), human myoblast cell line C2C12 (ATCC, CRL-1772), human cervical carcinoma cell line HelaRC32 (ATCC, CRL-2972), and human hepatoma HuH-7 cell line (National Collection of Authenticated Cell Cultures, SCSP-526) were cultured in Dulbecco’s Modified Eagle Medium (DMEM) (Thermo Fisher Scientific, C11995500BT) supplemented with 10% fetal bovine serum (FBS; Excell Bio, FSP500) and 1% penicillin-streptomycin (P/S; Biosharp, BL505A) and the hamster ovary cell line CHO-K1 (Cobioer Gene Technology Co., CBP60296) was maintained in F12K media (Sigma-Aldrich, N3520-10X1L) supplemented with 10% FBS and 1% P/S. All cells were maintained at 37 ℃ and 5% CO_2_.

### Virus packaging

To package AAV viruses for cell transduction and animal injection, HEK 293T cells were plated in 15 cm dishes at a density of 2E7 cells/dish. The next day, the Rep-Cap plasmid, pHelper plasmid, and ITR-containing plasmid AAV-CAG-Fluc-2A-eGFP were co-transfected into cells. At 72 h after transfection, cells were harvested by centrifugation at 500 × g for 10 min, resuspended in phosphate-buffered saline (PBS), and then the recombinant virus was released by freezing and thawing the cells three times. The crude lysate was clarified by centrifugation at 500 × g for 10 min and treated with benzonase at 250 U/mL final concentration at 37 °C for 30 min. Virus was further purified by iodixanol step gradient and heparin sulfate affinity chromatography. AAV titers were quantified by qPCR with a standard curve or by TaqMan-based droplet digital PCR (ddPCR), targeting the inverted terminal repeat (ITR) region. The viruses were stored at –80 °C at titers ranging from 1E12–1E13 genome copies/mL.

### AAV diameter

The particle size distribution of AAV preparations was assessed using dynamic light scattering (DLS) on the STUNNER™ nanoparticle analyzer (Unchained Labs, Pleasanton, CA, USA), following the manufacturer’s standard protocol. Prior to analysis, AAV samples were diluted in 1× PBS (pH 7.4) to an optimal concentration range (typically 1E12–1E13 vg/mL) to ensure accurate scattering signal and minimize multiple scattering effects. Measurements were conducted at 25 °C using disposable cuvettes, with each sample analyzed in triplicate to assess reproducibility. The hydrodynamic diameter and polydispersity index (PDI) were derived from the autocorrelation function of the scattered light intensity and analyzed using cumulant fitting models. Results were reported as mean hydrodynamic diameter (Z-average) and PDI, providing a quantitative evaluation of AAV particle size uniformity and aggregation status.

### RNA isolation and RT-qPCR

Total RNA was extracted from samples using the TransZol Up Plus RNA Kit (TransGen Biotech, ER501-01). The RNA concentration was determined using a Nanodrop (Thermo Fisher Scientific, 840-317400). Reverse transcription of 1 μg total RNA was performed with the EasyScript^®^ All-in-One First-Strand cDNA Synthesis SuperMix for qPCR (with One-Step gDNA Removal) kit (TransGen Biotech, AE341-03). Transgene expression was quantitated with 2× SYBR Green qPCR Master Mix (Bimake, B21203) following the manufacturer’s instructions. Procedures were run by StepOne Plus real-time PCR system (Applied Biosystems).

### Quantification of barcodes by next-generation sequencing

RNA was reverse transcribed into cDNA using the HiScript III 1st Strand cDNA Synthesis Kit (with gDNA wiper) (Vazyme, R312-02). PCR was performed with barcoded primers and the Q5 Hot Start High-Fidelity DNA Polymerase (NEB, M0493L). PCR amplicons were purified by Hieff NGS DNA Selection Beads (Yeason, 12601ES08) and sequenced using Illumina NovaSeq X Plus (Novogene).

### DNA isolation and quantification

Genomic DNA was extracted from tissue samples using the Hipure Universal DNA Kit (Magen Biotech, D3018-03) in accordance with the manufacturer’s protocol. Quantification of vector genome copies was carried out using ddPCR, targeting the firefly luciferase (*Fluc*) transgene and the endogenous *GAPDH* gene as a reference control gene (Fig. 1G and Supplementary Fig. 3B). ddPCR reactions were prepared using ddPCR Supermix for Probes (Bio-Rad, 1863024), with primers and a FAM-labeled probe specific to the *Fluc* sequence or *GAPDH* (Supplementary Table 3, synthesized by GENEWIZ). Droplet generation was performed using the Droplet Generator (Bio-Rad, QX200), followed by thermal cycling under standard conditions (95 °C for 10 min; 40 cycles of 94 °C for 30 s and 60 °C for 1 min; 98 °C for 10 min). Droplets were then read on the QX200 Droplet Reader, and data were analyzed with QuantaSoft software to determine absolute copy numbers. Absolute number of *Fluc* transgene and *GAPDH* molecules in each sample was quantified using a standard curve generated by amplifying the linearized CAG-Fluc-P2A-EGFP plasmid or a *GAPDH* containing sequence in each run by qPCR (Fig. 2E, 3D and Supplementary Fig. 3B).

### Western blot

Protein lysates from tissue samples were prepared in radioimmunoprecipitation assay (RIPA) buffer (Beyotime Biotechnology, P0013B) and centrifuged to remove debris. Total protein concentrations were determined using the modified BCA protocol (Sangon Biotech, C503051). Equal amounts of total protein from each sample were separated on 8–16% SurePAGE^TM^ precast polyacrylamide gels (GenScript, M00660). MOPS running buffer (GenScript, M00680-500) was used for electrophoresis. All gels were transferred to PVDF membranes in Western Rapid Transfer Buffer (Beyotime, P0572-2L). Membranes were blocked for 1 h in 5% skim milk (Beyotime, P0216-300g). All membranes were then incubated for 16 h at 4 ℃ with primary antibody diluted in primary antibody dilution buffer (Beyotime, P0023A-100mL). Primary antibodies used were against luciferase (1:2000, Proteintech, 27986-1-AP) and GAPDH (1:2000, Proteintech, 10494-1-AP). Blots were washed three times for 10 min each with Tris-buffered saline with Tween 20 (TBST) and then incubated with a goat anti-rabbit HRP-conjugated secondary antibody (Proteintech, SA00001-2) diluted 1:5000 in blocking buffer for 2 h at room temperature.

### Animal experiments

All relevant procedures involving animals were reviewed and approved by the Institutional Animal Care and Use Committee (IACUC) of South China Normal University and the Institutional Animal Care and Use Committee of Guangzhou Huazhen Bioscience (permit no. HZ2019027). Male mice (6–8 weeks old) were obtained from Zhuhai BesTest Bio-Tech Co., Ltd. and housed under a 12-hour light-dark cycle with *ad libitum* access to food and water. To evaluate the AAV transduction potential, mice were randomly assigned to experiments and transduced via intravenous injection (lateral tail vein) with a virus dose of 2E11 or 1E12 vg per mouse. Animals were euthanized 3 weeks after injection and tissues were harvested for the downstream transgene expression and biodistribution analyses. For NHP studies, 10 male 4–5 years old cynomolgus monkeys were purchased from Guangzhou Huazhen Biosciences Company Limited and housed in a controlled environment set to maintain a temperature of 18– 26 °C with a relative humidity of 60–80%. AAVs were intravenously administered on Day 0 and serum biochemistry were measured on Days 0, 1, 2, 3, 7, 14, and 28. Muscle biopsies were obtained on Day 14 and 28 under anesthesia using a muscle biopsy punch. Liver biopsies were obtained on Day 28 under anesthesia using a liver biopsy needle with guidance from ultrasound.

### In vivo imaging

Mice were injected with 150 mg/kg of D-luciferin (Promega, E1605) and anesthetized with isoflurane before imaging. The bioluminescence images were acquired 10 min after D-luciferin injection with a rate of one image per 2 min using the AniView100 multi-modality animal *in vivo* imaging system (Biolight Biotechnology). Total and average radiance was measured from the same size of the region of interest (ROI) using GV 6000ProII software (Biolight Biotechnology). The highest captured radiance over imaging time was determined as the peak of the kinetic curve and picked for analysis.

### Serum biochemistry assays

Each blood collection (about 1.0 mL per time point) was performed from peripheral vein of each animal into low binding tube containing 10 µL 0.5 M (K2) EDTA and placed on wet ice until centrifugation. Samples were centrifuged (3200 × g for 10 minutes at 2–8 °C) within one hour of collection. ALT, AST, and CK levels were measured using a clinical analyzer (Cobas e311, Roche).

### Immunofluorescence

Tissues harvested animals were fixed with 4% paraformaldehyde (PFA, Biosharp, XG1050) overnight at 4 ℃ and washed 3 × 10 min with Dulbecco’s PBS (DPBS). Fixed tissues were immersed in 30% sucrose at 4 ℃ until submersion, embedded in O.C.T. compound (Tissue-Tek, Biosharp, BL557A), and frozen at –80℃. Tissues were sectioned using FS800A cryostat (RWD Minux®) at a thickness of 10 μm. Sections were placed on poly-L-lysine coated slide glasses, dried for at least 30 minutes, and then stored at –20°C until being analyzed. Frozen sections were air-dried for 10 minutes at 37℃ prior to subsequent staining. For the microdystrophin immunostaining, tissue sections were incubated with a blocking solution consisting of PBS with 10% goat serum and 0.1% Triton X-100 for 30 minutes at room temperature (RT). Next, the sections were incubated with a rabbit polyclonal primary antibody against human dystrophin (1:200 in PBS; Abcam, ab15277) for 2 h at RT. After washing the sections 3 × 5 min with PBS, the sections were incubated with Alexa Fluor 555-labeled Donkey Anti-Rabbit IgG (H+L) secondary antibody (1:1000 in PBS; Beyotime, A0453) for an hour at RT. Nuclear counterstaining was performed with 4′,6-diamidino-2-phenylindole (DAPI). For the DAPI staining, tissue sections were washed 3 × 5 min with PBST (PBS + 0.1% Tween-20) and then mounted with anti-fluorescence quenched sealing solution (DAPI included) (Beyotime, P0131). Signals were detected with a confocal microscope (ZEISS LSM 800).

### Statistical analysis and data visualization

Data are presented as the mean ± standard error of the mean. Statistical analyses were carried out using GraphPad Prism Version 8 (GraphPad Software). An unpaired Student’s *t*-test was performed for comparison of two groups. Comparisons between three or more groups were analyzed with one-way ANOVA when there is only one independent variable or two-way ANOVA when there are two independent variables. A *p* value of 0.05 or less was considered significant in all experiments. Sequence alignments and schematic illustration of mice experiment were created with Biorender.

## Data availability statement

Source data for each relevant figure is provided in a Source Data file. The data that support the findings of this study are available from the corresponding author upon reasonable request.

## Acknowledgments

This work was supported by grants from Shenzhen Medical Research Fund (D2301004).

## Author contributions

Conceptualization: H.L., Y.B., Y.Z., and Y.P.; Investigation: Y.P., H.C., Y.Z., Z.D., J.C., K.T., X.C., D.Q., L.S., and X.T.; Methodology: Y.P., H.C., Y.Z., Z.D., L.S., and J.C.; Project administration: Y.B., Y.Z., and Y.P.; Resource: H.L., Y.B., M.G., R.D., Z.Z., and Z.Y.; Supervision: H.L., Y.B., and Y.P.; Validation: Y.P. and Y.B.; Visualization: Y.P. and H.C.; Original draft writing: Y.P.; Review and editing: G.G., H.L., Y.B., Y.P., Y.Z., Y.F., M.G., and R.D.

---

**Supplementary Figure 1.**
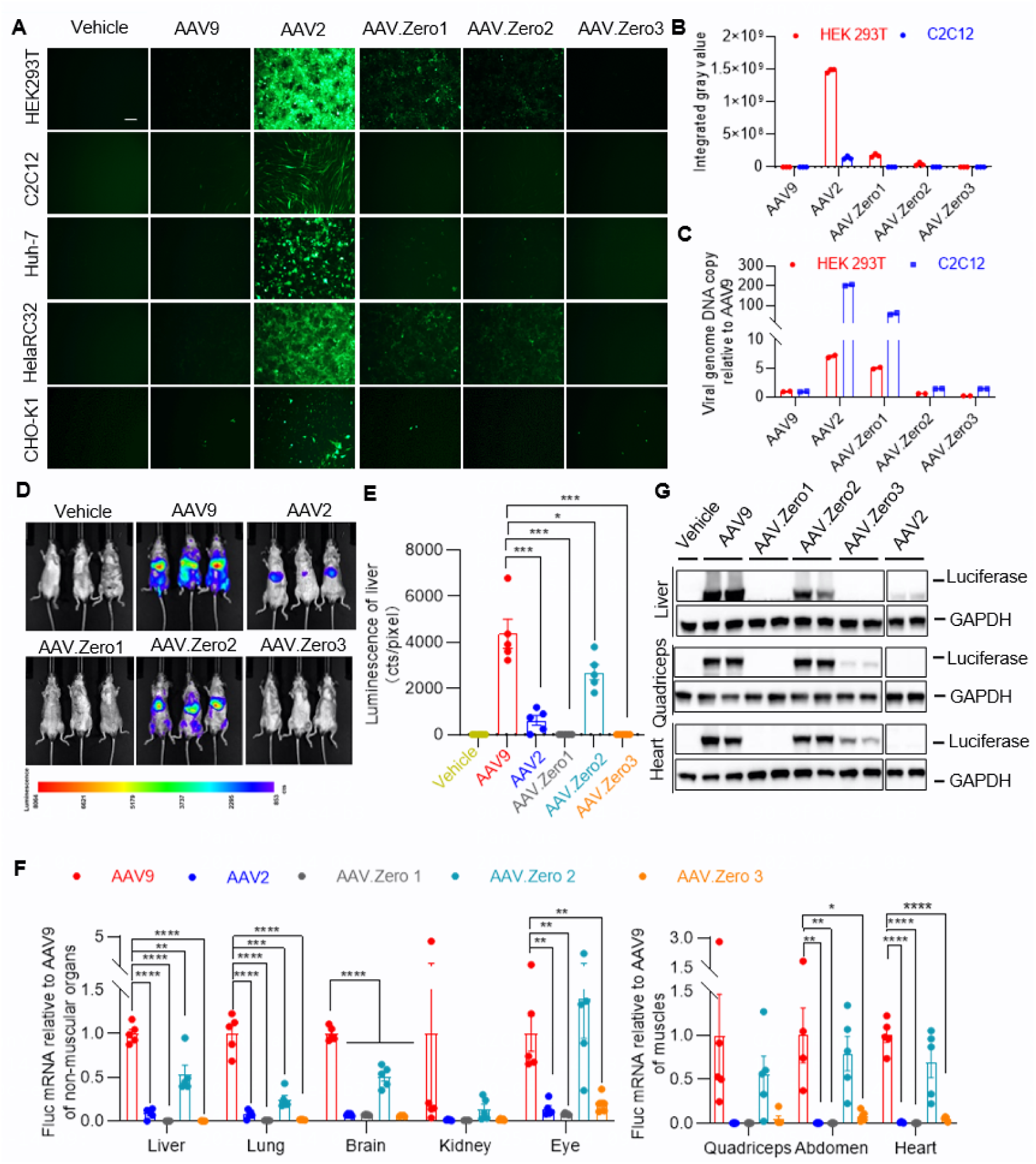
The tropisms of chimeric capsid variants systematically degenerate even at higher virus dose. A–C. HEK293T, C2C12, Huh-7, HelaRC32, and CHO-K1 cell lines being transduced with AAV9-, AAV2-, AAV.Zero1-, AAV.Zero2-, or AAV.Zero3-CAG-Fluc-P2A-EGFP at MOI=1E5 after 120 hours. A. Representative images of each treatment. Scale bar: 100 μm. B. Quantification of the integrated gray value of the EGFP fluorescence strength in HEK 293T and C2C12 cells (n=3). C. Quantification of relative virus genome copy numbers in HEK 293T and C2C12 cells (n=2). D–G. 8-week-old C57BL/6J mice were systemically injected with 1E12 vg per mouse (∼4E13 vg/kg) of AAV9-, AAV2-, AAV.Zero1-, AAV.Zero2-, or AAV.Zero3-CAG-Fluc-P2A-EGFP and data were collected 21 days post-injection. Representative whole body *in vivo* bioluminescence images (D). Quantification of firefly luciferase luminescence from liver (n=5) (E). Quantification of fold-difference in *Fluc* mRNA expression in various tissues (F) compared with normalized AAV9. The *p* values were calculated between AAV9 and each other groups by Student’s *t*-test (n=4–5), \**p* < 0.05, \*\**p* < 0.01, \*\*\**p* < 0.001 and \*\*\*\**p* < 0.0001. Representative western blot images detecting luciferase and GAPDH in quadriceps, heart, and liver (G).

**Supplementary Figure 2.**
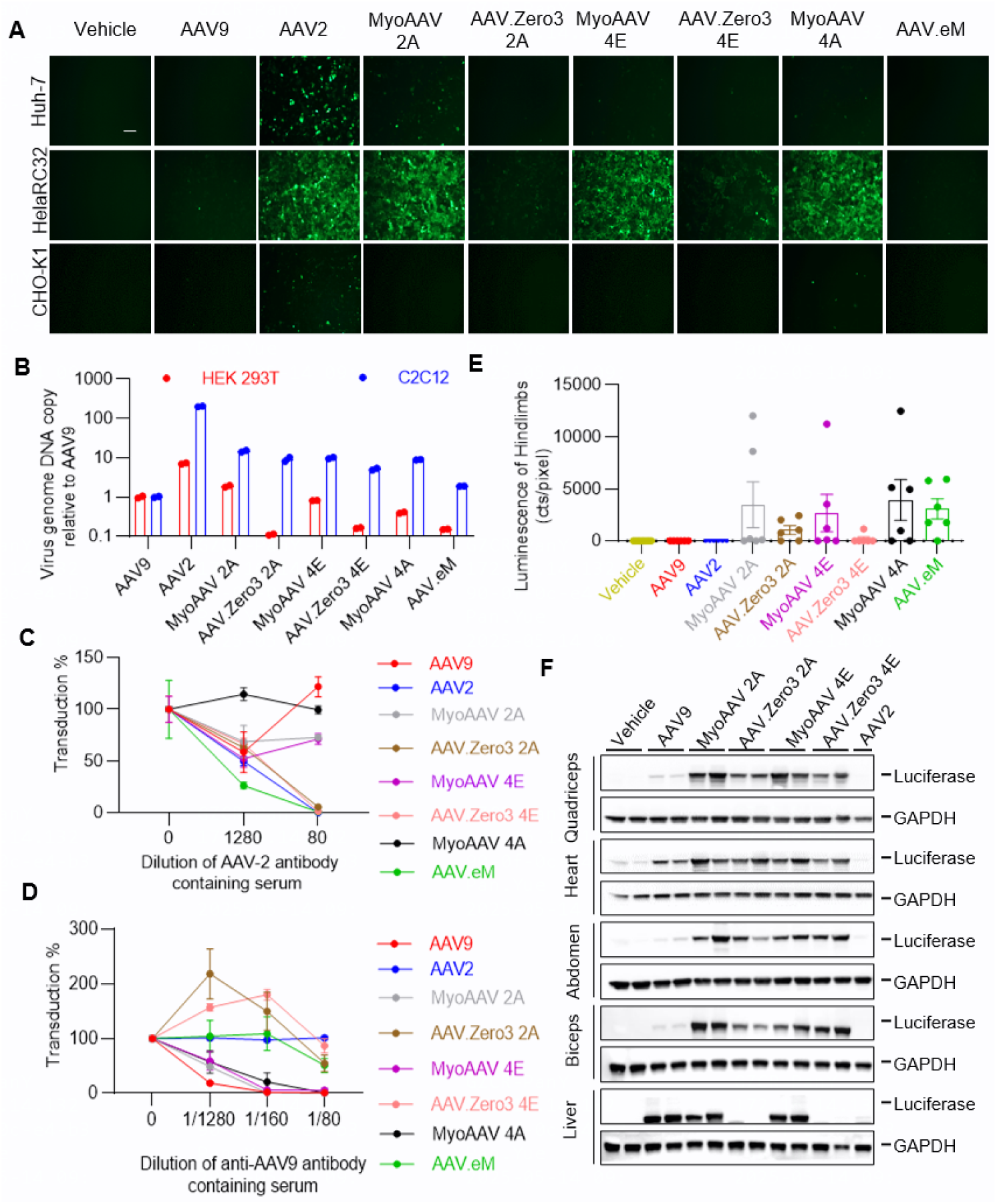
RGD-containing peptides enable chimeric capsid selectively target to muscle with high efficiency in C57BL/6J mice. A. Representative images Huh-7, HelaRC32, and CHO-K1 cell lines 72 hours after being transfected with AAV9-, AAV2-, MyoAAV 2A-, AAV.Zero3 2A-, MyoAAV 4E-, AAV.Zero3 4E-, MyoAAV 4A-, or AAV.eM-CAG-Fluc-P2A-EGFP at MOI=1E5. Scale bar: 100 μm. B. Quantification of relative virus genome copy numbers in HEK 293T and C2C12 cells transduced with AAV9-, AAV2-, MyoAAV 2A-, AAV.Zero3 2A-, MyoAAV 4E-, AAV.Zero3 4E-, MyoAAV 4A-, or AAV.eM-CAG-Fluc-P2A-EGFP at MOI=1E5 (n=2). C. Different dilutions of AAV2-antibody containing serum were incubated with constant amounts of virus at MOI=1E6 and tested for neutralization of transduction using an *in vitro* assay (n=3–4). D. Different dilutions of AAV9-antibody containing serum were incubated with constant amounts of virus at MOI=1E6 and tested for neutralization of transduction using an *in vitro* assay (n=3–6). E–F. 8-week-old C57BL/6J mice were systemically injected with 2E11 vg per mouse (∼8E12 vg/kg) of AAV9-, AAV2-, MyoAAV 2A-, AAV.Zero3 2A-, MyoAAV 4E-, AAV.Zero3 4E-, MyoAAV 4A-, or AAV.eM-CAG-Fluc-P2A-EGFP- and data were collected 21 days post-injection. Quantification of the firefly luciferase luminescence intensity from hindlimbs (n=6) (E). Representative western blot images detecting luciferase and GAPDH in quadriceps, heart, abdomen, biceps, and liver (F).

**Supplementary Figure 3.**
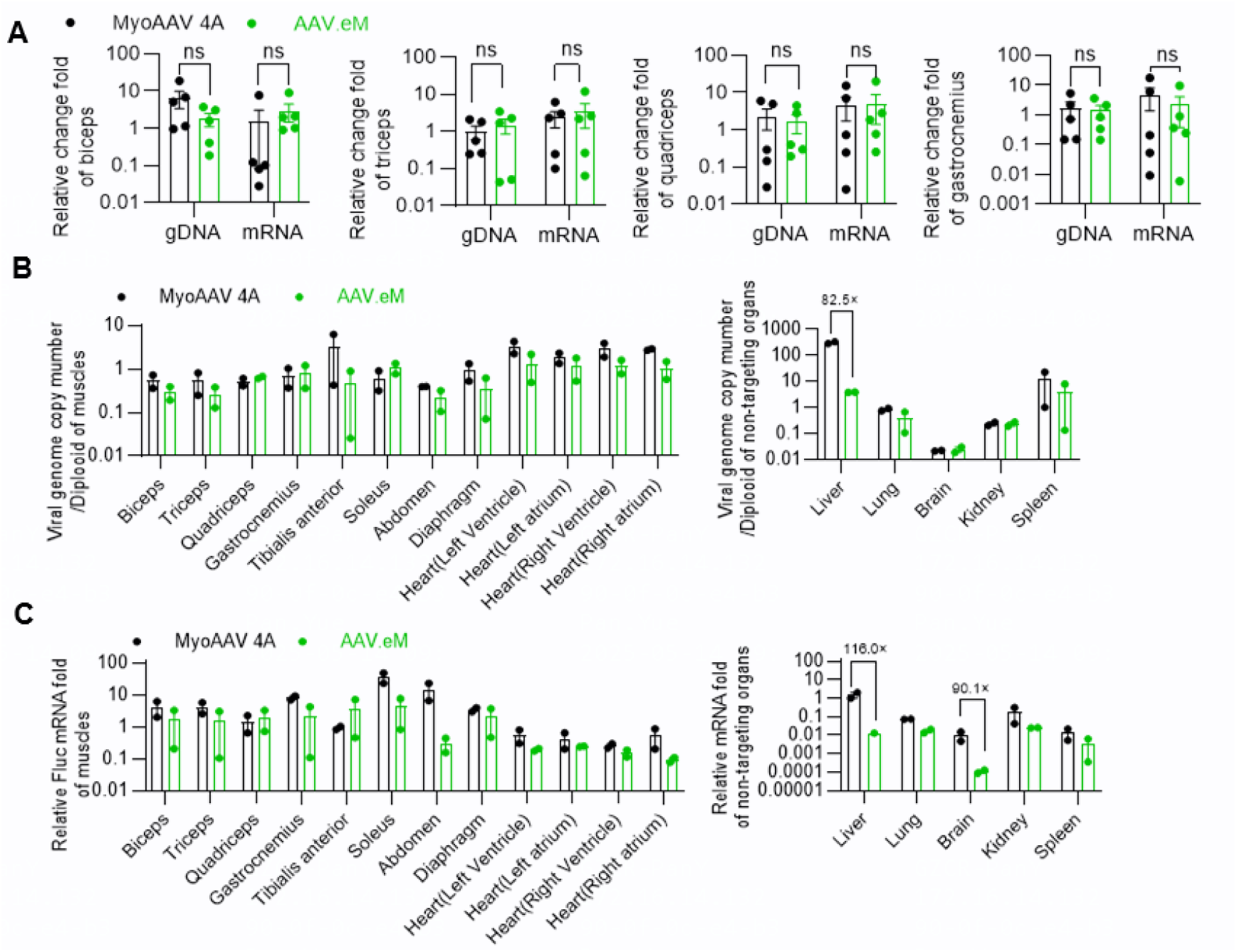
AAV.eM exhibits less off-target transduction and comparable muscle-specific targeting compared with MyoAAV 4A. A–C. NHPs were given intravenous administration of 3E13 vg/kg of MyoAAV 4A- or AAV.eM-CAG-Fluc-P2A-EGFP. Relative change fold of viral gDNA copy and *Fluc* mRNA in biceps, triceps, quadriceps and gastrocnemius sampled 28 days after virus injection (A). The *p* value was calculated by unpaired Student’s *t*-test (n=5), ns: not significant. The biceps, triceps, quadriceps, gastrocnemius, tibialis anterior, soleus, abdomen, diaphragm, heart, liver, lung, brain, kidney, and spleen were sampled 28 days after virus injection (n=2). Absolute quantification of viral vector genomes per diploid of each tissue was assayed by ddPCR (B). Fold-change in *Fluc* mRNA expression relative to *GAPDH* as assayed by qPCR with standard curve (C).

**Supplementary Figure 4.**
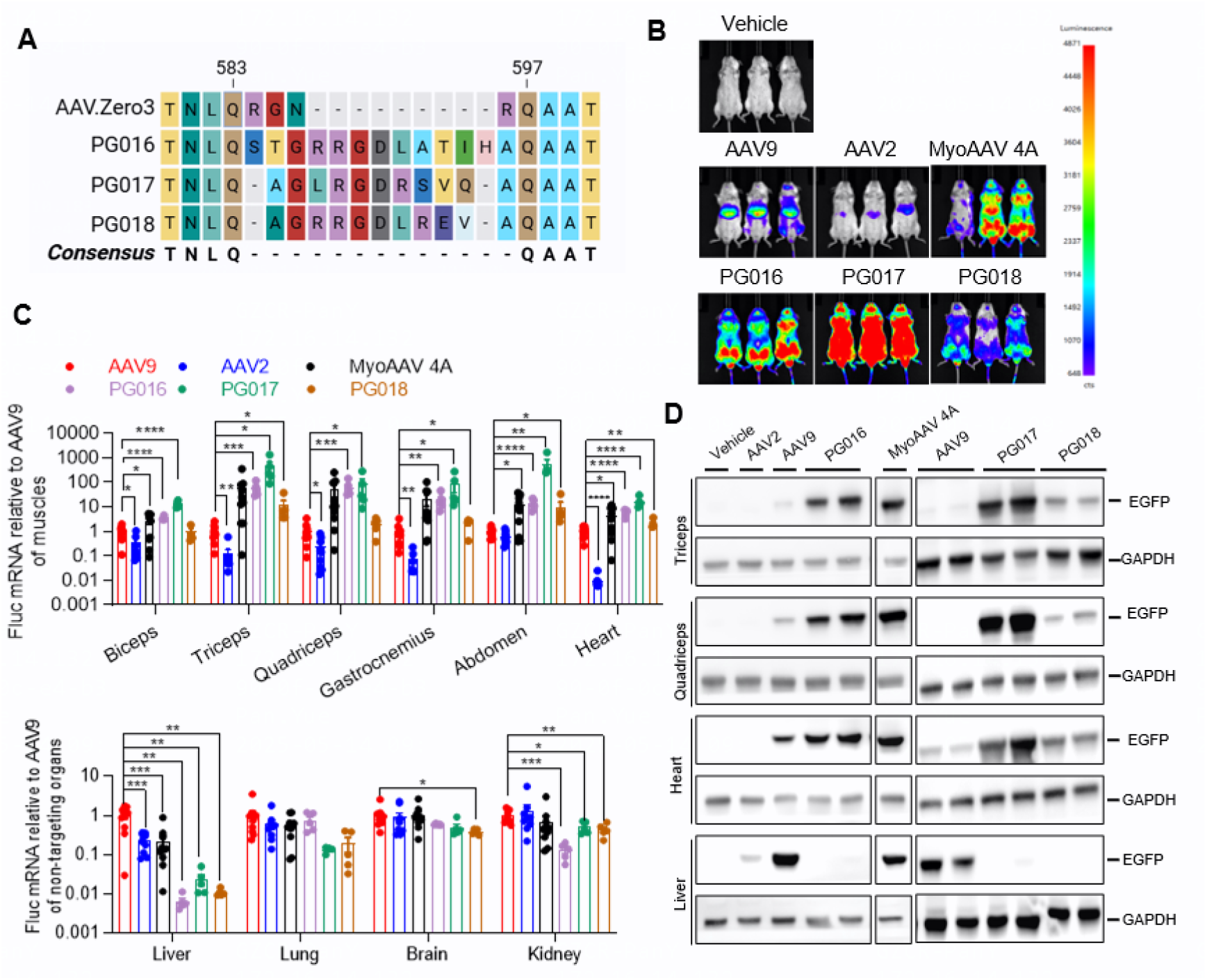
Novel muscle targeting peptides based on low-background capsid exhibit selectively target to muscles with high efficiency. A–D. 8-week-old BALB/c mice were systemically injected with 2E11 vg/mouse (∼8E12 vg/kg) of AAV9-, AAV2-, MyoAAV 4A, PG016, PG017, or PG018-CAG-Fluc-P2A-EGFP and tissues were collected 21 days post-injection. Sequence alignments of the RGD-containing peptide insertion region of PG016, PG017, and PG018 compared with AAV.Zero3 (A). Representative whole body *in vivo* bioluminescence images (B). Quantification of fold-difference in *Fluc* mRNA expression in various tissues (C) compared with normalized AAV9. The *p* values were calculated between AAV9 and each other group by Student’s *t*-test (n=9-10 except n=4–5 for PG016, PG017 and PG018). Representative western blot images detecting EGFP and GAPDHs in varied tissues (D).

